# Cuttlefish: Fast, parallel, and low-memory compaction of de Bruijn graphs from large-scale genome collections

**DOI:** 10.1101/2020.10.21.349605

**Authors:** Jamshed Khan, Rob Patro

## Abstract

**Motivation:** The construction of the compacted de Bruijn graph from collections of reference genomes is a task of increasing interest in genomic analyses. These graphs are increasingly used as sequence indices for short and long read alignment. Also, as we sequence and assemble a greater diversity of genomes, the colored compacted de Bruijn graph is being used as the basis for efficient methods to perform comparative genomic analyses on these genomes. Therefore, designing time and memory efficient algorithms for the construction of this graph from reference sequences is an important problem.

**Results:** We introduce a new algorithm, implemented in the toolCuttlefish, to construct the (colored) compacted de Bruijn graph from a collection of one or more genome references. Cuttlefish introduces a novel approach of modeling de Bruijn graph vertices as finite-state automata; it constrains these automata’s state-space to enable tracking their transitioning states with very low memory usage. Cuttlefish is fast and highly parallelizable. Experimental results demonstrate that it scales much better than existing approaches, especially as the number and the scale of the input references grow. On our test hardware, Cuttlefish constructed the graph for 100 human genomes in under 9 hours, using ~29 GB of memory while no other tested tool completed this task. On 11 diverse conifer genomes, the compacted graph was constructed by Cuttlefish in under 9 hours, using ~84 GB of memory, while the only other tested tool that completed this construction on our hardware took over 16 hours and ~289 GB of memory.

**Availability:** Cuttlefish is written in `C++14`, and is available under an open source license at https://github.com/COMBINE-lab/cuttlefish.

**Supplementary information:** Supplementary text are available at *Bioinformatics* online.

## 1 Introduction

With the increasing affordability and throughput of sequencing, the set of whole genome references available for comparative analyses and indexing has been growing tremendously. Modern sequencing technologies can generate billions of short read sequences per-sample, and with state-of-the-art *de novo* and reference-guided assembly algorithms, we now have thousands of mammalian-sized genomes available. Moreover, we have now successfully assembled genomes that are an order of magnitude larger than typical mammalian genomes, with the largest ones among these being the sugar pine (~31 Gbp) (Stevens *et al.*, 2016) and the Mexican walking fish (~32 Gbp) (Nowoshilow *et al.*, 2018). With modern long-read sequencing technologies and assembly techniques, the diversity and the completeness of reference-quality genomes is expected to continue increasing rapidly. Representation of these reference collections in compact forms facilitating common genomic analyses are thus of acute interest and importance, as are efficient algorithms for constructing these representations.

To this end, the de Bruijn graph has become an object of central importance in many genomic analysis tasks. While it was initially used mostly in the context of genome (and transcriptome) assembly (EULER (Pevzner *et al.*, 2001), EULER-SR (Chaisson and Pevzner, 2008), Velvet (Zerbino and Birney, 2008; Zerbino *et al.*, 2009), ALLPATHS (Butler *et al.*, 2008; MacCallum *et al.*, 2009), ABySS (Simpson *et al.*, 2009), Trans-AByss (Robertson *et al.*, 2010), SPAdes (Bankevich *et al.*, 2012), Minia (Chikhi and Rizk, 2013), SOAPdenovo (Li *et al.*, 2010; Luo et al., 2015)), it has seen increasing use in comparative genomics (Cortex (Iqbal *et al.*, 2012), DISCOSNP (Uricaru *et al.*, 2014), Scalpel (Fang *et al.*, 2016), BubbZ(Minkin and Medvedev, 2020)), and has also been used increasingly in the context of indexing genomic data, either from raw sequencing reads (Vari (Muggli *et al.*, 2017), Mantis (Pandey *et al.*, 2018; Almodaresi *et al.*, 2019), VariMerge (Muggli *et al.*, 2019), MetaGraph (Karasikov *et al.*, 2020)), or from assembled reference sequences (deBGA (Liu *et al.*, 2016), Pufferfish (Almodaresi *et al.*, 2018), deSALT (Liu *et al.*, 2019)), or from both (BLight (Marchet *et al.*, 2019), Bifrost (Holley and Melsted, 2020)). These latter applications most frequently make use of the (colored) compacted deBruijn graph, a variant of the de Bruijn graph in which the maximal non-branching paths (also referred to as unitigs) are condensed into single vertices in the underlying graph structure. This retains all the information of the original graph, while typically requiring much less space to store, index, and process. The set of maximal unitigs for a de Bruijn graph is unique and forms a node decomposition of the graph (Chikhi *et al.*, 2016). The colored variant is built over multiple references.

The (colored) compacted de Bruijn graph has become an increasingly useful data structure in computational genomics, for its use in detecting and representing variation within a population of genomes, comparing whole genome sequences, as well as, more recently, its accelerated use as a foundational structure to assist with indexing one or more related biological sequences. Applying de Bruijn graphs for these whole genome analysis tasks highlights a collection of algorithmic challenges, such as efficient construction of the graphs (out-of-core (Kundeti *et al.*, 2010), khmer (Pell *et al.*, 2012), BOSS (Bowe *et al.*, 2012), Minia (Chikhi and Rizk, 2013), kFM-index (Rødland, 2013), DBGFM (Chikhi *et al.*, 2014), BFT (Holley *et al.*, 2016)), memory-efficient representations for the graphs once constructed (SplitMEM (Marcus *et al.*, 2014), BWT-based algorithm (Baier *et al.*, 2015), TwoPaCo (Minkin *et al.*, 2016), deGSM (Guo *et al.*, 2019), Bifrost (Holley and Melsted, 2020)), building space- and time-efficient indices on these representations (BFT (Holley *et al.*, 2016), Pufferfish (Almodaresi *et al.*, 2018), BLight (Marchet *et al.*, 2019)), and performing read-alignment on the graphs using the indices (deBGA (Liu *et al.*, 2016), deSALT (Liu *et al.*, 2019), PuffAligner (Almodaresi *et al.*, 2020)), to mention a few.

In this work, we tackle the initial step of the pipelines associated with whole genome and pan-genome analysis tasks based on de Bruijn graphs: the time- and memory-efficient construction of (colored) compacted de Bruijn graphs. We present a novel algorithm, implemented in the tool Cuttlefish, for this purpose. It has excellent scaling behavior and low memory requirements, exhibiting better performance than state-of-the-art tools for compacting de Bruijn graphs from reference collections. The algorithm models each distinct k-mer (i.e. vertex of the de Bruijn graph) of the input reference collection as a finite-state automaton, and designs a compact hash table structure to store succinct encodings of the states of the automata. It characterizes the k-mers that flank maximal unitigs through an implicit traversal over the original graph — without building it explicitly — and dynamically transitions the states of the automata with the local information obtained along the way. The maximal unitigs are then extracted through another implicit traversal, using the obtained k-mer characterizations.

We assess the algorithm on both individual genomes, and collections of genomes with diverse structural characteristics: 7 closely related humans; 7 related but different apes; 11 related, different, and huge conifer plants; and 100 humans. We compare our tool to three existing state-of-the-art tools: Bifrost, deGSM, and TwoPaCo. Cuttlefish is competitive with or faster than these tools under various settings of k-mer length and thread count, and uses less memory (often multiple *times* less) on all but the smallest datasets.

## 2 Related work

The amount of short-read sequences produced from samples, like whole genome DNA, RNA, or metagenomic samples, can be in the order of billions. Construction of long-enough contiguous sequences, also referred to as contigs (Staden, 1980), from the sets of reads is known as the fragment assembly problem, and is a central and long-standing problem in computational biology. Many short-read fragment assembly algorithms (BCALM (Chikhi *et al.*, 2014), BCALM2 (Chikhi *et al.*, 2016), Bruno (Pan *et al.*, 2018), deGSM (Guo *et al.*, 2019)) use the de Bruijn graph to represent the input set of reads, and assemble fragments through graph compaction. More generally, fragment assembly algorithms are typically present as parts of larger and more complex genome assembly algorithms. In this work, we study the closely related problem of constructing and compacting de Bruijn graphs in the whole genome setting. While the computational requirements for the compacted de Bruijn graph construction from genome references are typically substantial compared to the latter phases of sequence analysis (whole- and pan-genome) tasks, only a few tools have focused on the specific problem.

SplitMEM (Marcus *et al.*, 2014) exploits topological relationships between suffix trees and compacted de Bruijn graphs. It introduces a construct called *suffix skips*, which is a generalization of suffix links and is similar to pointer jumping techniques, to fast navigate suffix trees for identifying MEMs (maximal exact matches), and then transforms those into vertices and edges of the output graph. SplitMEM is improved upon using two algorithms (Baier *et al.*, 2015): based on a compressed suffix tree, and on the Burrows-Wheeler Transform (BWT) (Burrows and Wheeler, 1994).

TwoPaCo (Minkin *et al.*, 2016) takes the approach of enumerating the edges of the original de Bruijn graph, which aids in identifying the *junction* positions in the genomes, i.e. positions that correspond to vertices in TwoPaCo’s compacted graph format. Initially having the entire set of vertices as junction candidates, it shrinks the candidates set using a Bloom filter (Bloom, 1970), with a pass over the graph. Since the Bloom filter may contain false positive junctions, TwoPaCo makes another pass over the graph, this time keeping a hash table for the reduced set of candidates and thus filtering out the false positives.

deGSM (Guo *et al.*, 2019) builds a BWT of the maximal unitigs without explicitly constructing the unitigs themselves. It partitions the (k + 2)-mers of the input into a collection of buckets, and applies parallel external sorting on the buckets. Then characterizing the k-mers at the ends of the maximal unitigs using the sorted information, it merges the buckets in a way to produce the unitigs-BWT.

Bifrost (Holley and Melsted, 2020) initially builds an approximate de Bruijn graph using blocked Bloom filters. Then for each k-mer in the input, it extracts the maximal unitig that contains that k-mer, by extending the k-mer forward (and backward), constructing the suffix (and prefix) of the unitig. This produces an approximate compacted de Bruijn graph. Then using a k-mer counting and minimizer-based policy, it refines the extracted unitigs by removing the false positive k-mers present in the compacted graph.

## 3 Preliminaries

We consider all strings to be over the alphabet Σ *=* {*A, C, G, T*}. For some string s, |s| denotes its length. *s*[*i..j*] denotes the substring of s from the *i*’th to the *j*‘th character, inclusive (with 1-based indexing). *suf_ℓ_*(*s*) and *pre_ℓ_*(*s*) denote the suffix and the prefix of *s* with length *ℓ*, respectively. For two strings *x* and *y* such that *suf_ℓ_*(*x*) = *pre_ℓ_*(*y*), the *glue* operation ⊙ is defined as *x*⊙^*ℓ*^*y* = *x·y* [*ℓ*+1..|*y*|], where (·) is the append operation.

A *k-mer* is a string of length *k*. For some string *x*, its *reverse complement* 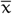 is the string obtained by reversing the order of the characters in x and complementing each character according to the nucleotide bases’ complementarity. The *canonical form* 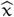 of a string *x* is the string 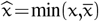, according to the lexicographic ordering.

For a set *S* of strings and an integer *k*>0, the corresponding *de Bruijn graph G*(*S,k*) is defined as a directed graph with: (i) its set of vertices being exactly the set of *k*-mers of *S*; and (ii) two vertices *u* and *v* being connected with a directed edge *u*→*v* iff there exists some (*k*+1)-mer *e* in *S* such that *pre_k_*(*e*) = *u* and *suf_k_*(*e*) = *v*. Note that, we adopt an *edge-centric* definition of the de Bruijn graph where an edge exists iff there is some corresponding (*k*+1)-mer present in *S*, as opposed to the *node-centric* definition where the edges are implicit given the vertices, i.e. an edge *u*→*v* exists iff *suf*_*k*−1_(*u*) = *pre*_*k*−1_(*v*). A *walk* in *G*(*S,k*) is a sequence *w* = (*v*_0_,…, *v_m_*) of vertices such that for any two successive vertices *v_i_* and *v*_*i*+1_ in *w*, there exists an edge *v_i_* → *v*_*i*+1_ in *G*(*S,k*). A *path* is a walk with no repeated vertex. A path *p* = (*v*_0_,…, *v_m_*) is said to be a *unitig* if |*p*| = 1, or for all 0<*i*<*m*, the in- and out-degrees of *v_i_* are 1, and the out-degree of vo and the in-degree of *v_m_* are 1.

For our purposes, we use the bidirected variant of de Bruijn graphs. For the bidirected de Bruijn graph *G*(*S,k*), the set of vertices is exactly the set of canonical *k*-mers obtained from *S*. A vertex *v* is said to have exactly two *sides*, referred to as the *front* side *s*_*v*f_ and the *back* side *s*_*v*b_. An edge exists between two vertices *u* and *v* iff there is some (*k*+1)-mer *e* in *S* such that *u* and *v* are the canonical forms of the *k*-mers *pre_k_*(*e*) and *suf_k_*(*e*), respectively; i.e. 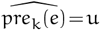 and 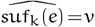. *e* is incident to exactly one side of *u* and one side of *v*. These incidence sides are determined using the following rule:

1. If *pre_k_*(*e*) = *u*, i.e. *pre_k_*(*e*) is in its canonical form, then *e* is incident to the back side *s*_*u*b_ of *u*; otherwise it is incident to *u*’s front side *s*_*u*f_.
2. If *suf_k_*(*e*) = *v*, i.e. *suf_k_*(*e*) is in its canonical form, then *e* is incident to the front side *s*_*v*f_ of *v*; otherwise it is incident to *v*’s back side *s*_*v*b_.

If *u=v*, then e is said to be a loop. So, an edge *e* can be defined as a 4-tuple (*u,s_u_,v,s_v_*), with *s_u_* and *s_v_* being the sides of the vertices *u* and *v* respectively to which *e* is incident to. The canonical *k*-mer 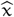 corresponding to a vertex *v* is referred to as its *label*, i.e. 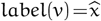. An example illustration of a bidirected de Bruijn graph is given in Fig. 1a.

**Fig. 1:**
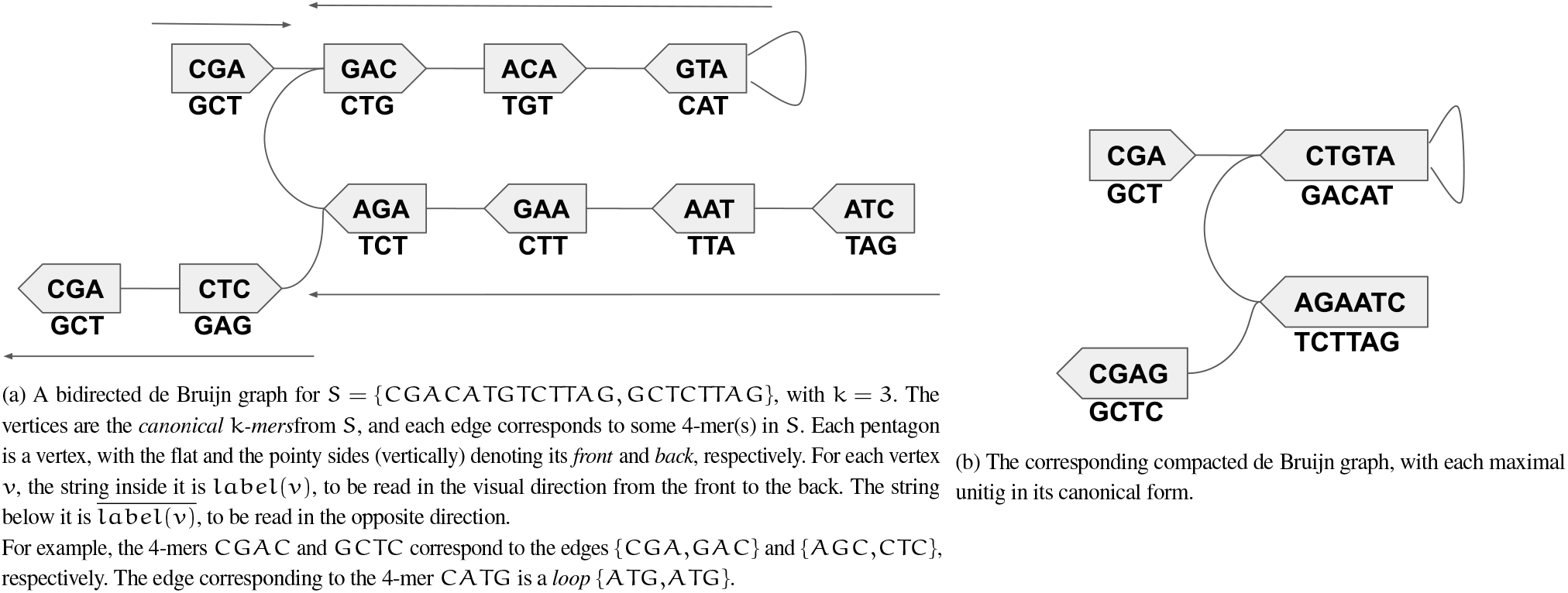
For *G*(*S,k*) in Fig. (1a), *w*_1_ = (*GAC,ACA,ATG,ATG*) (edges not listed) is a *walk*. *w*_1_ *spells* the string *GACATG*. It is not a *path* as the vertex *ATG* is repeated here. Whereas the walk *w*_2_ = (*GAC,ACA,ATG*) is a path, spelling *GACAT*. Besides, it is a *unitig*, and also a maximal one as it cannot be extended on either side while retaining itself a unitig. There are four *maximal unitigs* in the graph (the paths referred with the arrows), with the canonical spellings: *CGA, ATGTC, CTAAGA*, and *GAGC*.

A walk is defined as an alternating sequence of vertices and edges, *w* = (*v*_0_, *e*_1_, *v*_1_,…, *v*_*m*−1_, *e_m_, v_m_*), such that any two successive vertices *v*_*i*−1_ and *v*, in *w* are connected with the edge *e*, (through any side). *v*_0_ and *v_m_* are called the *endpoints*, and the *v*, for 0<*i*<*m* are called the *internal vertices* of the walk. *w* is said to *enter v_i_*, using *e_i_*, and *exit v*, using *e*_*i*+1_. For the endpoints *v*_0_ and *v_m_, w* does not have an entering and an exiting edge, respectively. For 0<*i* ≤ *m, w* is said to enter *v_i_* through its front side *s*_*v*_*i* f__ if *e_i_* is incident to *s*_*v*_*i*f__. Otherwise, it is said to enter *v_i_* through its back side *s*_*v*_*i* b__. For 0 ≤ *i*<*m*, the terminology for the exiting side is similarly defined using the incidence side of *e*_*i*+1_ to *v_i_*.

*w* is said to be *input-consistent* if, for each of its internal vertices *v_i_* the sides of *v*, to which *e*, and *e*_*i*+1_ are incident are different. Intuitively, an input-consistent walk enters and exits a vertex through different sides. We prove in lemma 1 (see Supplementary Sec. 2) that the reconstruction (spelling) of the input strings *s* ∈ *S* are possible only from input-consistent walks over *G*(*S,k*). Therefore, we are only interested in such walks, and refer to input-consistent walks whenever using the term walk onward.

For a vertex *v_i_* in *w*, let its label be *l_i_*, i.e. *l_i_* = label(*v_i_*). We say that *w* sees the *spelling s_i_* for the vertex *v_i_* (for 0<*i* ≤ *m*), such that *s_i_* is *l_i_* if *w* enters *v*, from the front, and is 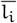 otherwise. *s*_0_ is defined analogously but using the exiting edge *e*_1_ for *v*_0_. The *spelling* of *w* is defined as 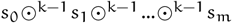.

A path *p* = (*v*_0_, *e*_1_, *v*_1_,…, *e_m_, v_m_*) is said to be a *unitig* if |*p*| = 1, or in *G*(*S,k*):

1. each internal vertex *v_i_* has exactly two incident edges, *e_i_* and *e*_*i*+1_;
2. and *v*_0_ has exactly one edge *e*_1_ and *v_m_* has exactly one edge *e_m_* incident to the sides respectively through which *p* exits *v*_0_ and enters *v_m_*.

A unitig is said to be maximal if it cannot be extended by a vertex on either side. Fig. 1 illustrates examples of walks, paths, their spellings, and maximal unitigs in bidirected de Bruijn graphs.

The *compacted de Bruijn graph G_c_*(*S,k*) of the graph *G*(*S,k*) is obtained by collapsing each of its maximal unitigs into a single vertex. Fig. 1b shows the compacted form *G_c_*(*S,k*) of the graph *G*(*S,k*) from Fig. 1a. The problem of compacting a de Bruijn graph is to compute the set of its maximal unitigs.

We use a slightly modified definition for the de Bruijn graph *G*(*S,k*) in solving this problem. We refer a vertex *v* as a *sentinel* if the first or the last *k*-mer *x* of some input string *s* ∈ *S* corresponds to *v*. We remove any edge incident to the side of *v* that is empty at the sentinel-inducing occurrence of *x*. This is to ensure correct reconstruction-ability of the input strings in *S* from *G_c_* (*S,k*) — so that each *s* ∈ *S* can be expressed as a sequence tiling of complete maximal unitigs. For example, including a new string *ACGA* to *S* at Fig. 1a should introduce the edge {*ACG,CGA*}. But we do not allow *CGA* to have an edge incident to its front side, since this side is empty at the sentinel-inducing occurrence of *CGA* as the first *k*-mer in the first input string — so this edge is not added to the graph.

## 4 Algorithm

### 4.1 Motivation

Given a set S of strings and an odd integer *k* > 0, a simple naive algorithm to construct the corresponding compacted de Bruijn graph *G_c_*(*S,k*) is to first construct the ordinary de Bruijn graph *G*(*S,k*), and then enumerate all the maximal non-branching paths from *G*(*S,k*). The graph construction can be performed by a linear scan over each *s* ∈ *S* storing information along the way in some graph representation format, like an adjacency list. Enumeration of the non-branching paths can be obtained using a linear-time graph traversal algorithm (Cormen *et al.*, 2009). However, storing the ordinary graph *G*(*S,k*) usually takes an enormous space for large genomes, and there is no simple way to traverse it without having the entire graph present in memory. This motivates the design of methods able to build *G_c_*(*S,k*) directly from *S* without having to construct *G*(*S,k*).

For some *s* ∈ *S*, it is important to note that by the definition of edge-centric de Bruijn graphs, each (*k*+1)-mer in s is an edge of *G*(*s,k*). Therefore, we can obtain a complete walk traversal *w*(*s*) over *G*(*s,k*) through a linear scan over *s*, without having to build *G* (*s,k*) explicitly. Also, each maximal unitig of the graph is contained as a subpath in this walk, as proven in lemma 2 (see Supplementary Sec. 2). For the set of strings *S*, the maximal unitigs are similarly contained in a collection of walks *W*(*S*).

Thus, to construct *G_c_*(*S,k*) efficiently, one approach is to extract off the maximal unitigs from these walks *W*(*S*), without building *G*(*S,k*). We describe below an asymptotically and practically efficient algorithm that performs this task.

### 4.2 Flanking vertices of the maximal unitigs

Similar to the TwoPaCo (Minkin *et al.*, 2016) algorithm, our algorithm is based on the observation that there exists abijection between the maximal unitigs of *G*(*S,k*) and the substrings of the input strings in *S* whose terminal k-mers correspond to the endpoints of the maximal unitigs. We refer to these endpoint vertices as *flanking* vertices. This observation reduces the graph compaction problem to the problem of determining the set of the flanking vertices. Once each vertex in *G*(*S,k*) can be characterized as either flanking or internal with respect to the maximal unitig containing it, the unitigs themselves can then be enumerated using a walk over *G*(*S,k*), by identifying subpaths having flanking *k*-mers at both ends.

Consider a maximal unitig *p* and one of its endpoints *v*. Say that *v* is connected to *p* through its side *s*_*v*_in__, and its other side is *s*_*v*_out__. *v* is a flanking vertex as it is not possible to extend p through *s*_*v*_out__ while retaining itself a unitig, due to one of the following cases:

i. there are either zero or multiple edges incident to *s*_*v*_out__; or
ii. there is exactly one edge (*v,s*_*v*_out__, *u,s_u_*) incident to *s*_*v*_out__, but *s*_*u*_ has multiple incident edges.

For trivial unitigs^1^, extending the unitig in both the scenarios violates the second property of (non-trivial) unitigs; while for non-trivial unitigs, the first property is violated in the first scenario and the other one is violated in the second scenario.

From this, we can observe that the adjacency information of the sides of the vertices can be used to determine the flanking vertices. As per lemma 4 (see Supplementary Sec. 2), a side of a vertex can have at most four distinct edges. This being finite, the adjacency information can be tracked using some data structure. Through a set of walks over *G*(*S,k*), each encountered edge (*u,s_u_,v,s_v_*) can be recorded for the sides *s_u_* and *s_v_*. With another set of walks, the maximal unitigs can then be extracted as per the flanking vertices, determined using the obtained adjacency information.

Another approach — adopted in TwoPaCo (Minkin *et al.*, 2016)) — is to have an edge list data structure supporting queries, and filling it up with the graph edges (i.e. (*k*+1)-mers) with a set of walks. In another set of walks, the flanking vertices can be characterized using incidence queries (four per side), and the maximal unitigs can be extracted on the fly.

### 4.3 A deterministic finite-state automaton model for vertices

In these characterization approaches of the flanking vertices, a source of redundancy in the information stored is the vertex sides that have multiple incident edges, or are sentinels (so, blocked off). By definition, these sides belong to the flanking vertices of the maximal unitigs. So, in a first set of walks over the graph, once a side of a vertex has been determined to have more than one distinct edge, or to be a sentinel side, the adjacency information for the side becomes redundant: for our purposes, we do not need to be able to enumerate the incident edges for this side, or to differentiate between them. Only the information that this side makes the vertex flanking is sufficient. Complementarily, another source of redundant information is the vertex sides observed to have exactly one incident edge up-to some point in the walks. For such a side, keeping track of only that single edge is sufficient, instead of having options to track more distinct edges: when a different edge is encountered, distinguishing between the edges becomes redundant.

Thus we observe that the only required information to keep per side of a vertex is:

i. whether it has exactly one distinct edge, and if so, which edge it is among the four possibilities (as per lemma 4, see Supplementary Sec. 2);
ii. or if there are multiple (or zero) distinct incident edges.

Hence, a side can have one of five different configurations: one for each possible singleton edge, and one for when it does not have exactly one distinct edge. This implies that each vertex can be in one of (5 × 5)= 25 different configurations. We refer to such a configuration as a *state* for a vertex. Figuring out the states of the vertices provides us with enough information to extract the maximal unitigs.

To compute the actual state of each vertex *v* in *G*(*S,k*), we treat *v* as a deterministic finite-state automaton (DFA). Prior to traversing *G*(*S,k*), no information is available for *v* — we designate a special state for such, the *unvisited* state. Then in response to each incident edge or sentinel *k*-mer visited for *v*, an appropriate state-transition is made.

Formally, a vertex *v* in *G*(*S,k*) is treated as aDFA (*Q*, Σ′, *δ, q*_0_, *Q′*), where —

#### States

*Q* = *Q′* ≪{*q*_0_} is the set of all possible 26 states. The 25 actual states (configurations) of the vertices in *G*(*S,k*) can be partitioned into four disjoint classes based on their properties, as in Fig. 2a.

**Fig. 2:**
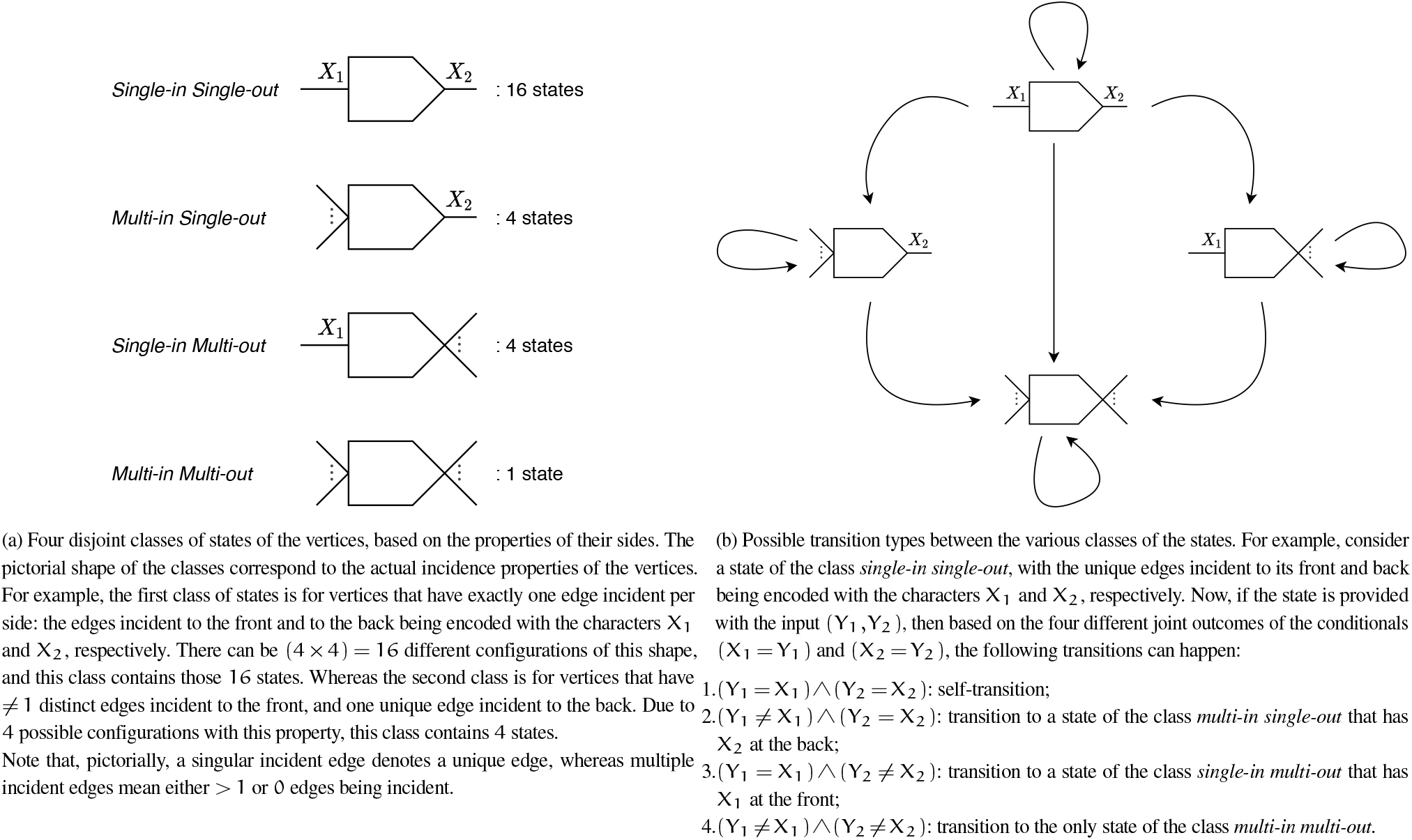
Classes of states of the vertices, and the transition relationships among those.

#### Input symbols

Σ′ = {(*c*_1_, *c*_2_)| *c*_1_, *c*_2_ ∈ Σ∪{ϵ}} (where ϵ denotes the empty character, used for sentinel-sides) is the set of input symbols for the automaton. In the algorithm, we actually make state-transitions not per edge, but per vertex. That is, for some walk *w* = (*v*_0_,…, *e_i_,v_i_,e*_*i*+1_,…, *v_m_*), we process the vertices *v_i_*. Processing *v_i_* means checking the edges *e_i_* and *e*_*i*+1_ simultaneously and only for *v_i_* (excluding the edges’ other endpoints); making state transitions for *v_i_*’s automaton as required^2^. Thus, the two incident edges {*e_i_,e*_*i*+1_,} are being used as input for the automaton of *v_i_*. The edges can be represented succinctly with a pair of characters (*c*_1_, *c*_2_) for *v_i_. c*_1_ and *c*_2_ encode the edges incident to *v_i_*’s front and back respectively, in the subwalk (*e_i_,v_i_,e*_*i*+1_)^3^.

#### Transition function

*δ*: *Q* × Σ′ → *Q* is the function governing the state-transitions. Fig. 2b presents a high-level view of the possible types of transitions between states of the four classes. Supplementary Fig. 1 illustrates a detailed overview of the transitions defined for the states as per various input symbols.

#### Initial state

*q*_0_ is the initial state for the automaton, which is the unvisited state.

#### Accept states

*Q′* is the set of the possible 25 different configurations (i.e. states) for the vertices.^4^

### 4.4 The Cuttlefish algorithm

For a set *S* of input strings, the proposed algorithm, Cuttlefish(*S*), works briefly as follows. Cuttlefish treats each vertex of the de Bruijn graph *G*(*S,k*) as a DFA — based on the novel modeling scheme introduced in subsection 4.3. First, it enumerates the set *K* of vertices of *G*(*S,k*), applying a *k*-mer counting algorithm on *S*. Then it builds a compact hash table structure for *K* — employing a minimal perfect hash function—to associate the automata to their states (yet to be determined). Having instantiated a DFA *M_v_* for each vertex *v* ∈ *K*, with *M_v_* being in the initial state *q*_0_ (i.e. the unvisited state), it makes an implicit traversal W(*S*) over *G*(*S,k*). For each instance (i.e. *k*-mer) *x* of each vertex *v* visited along *W*(*S*), it makes an appropriate state transition for *v*’s automaton *M_v_*—using the local information obtained for *x*, i.e. the two incident edges of *x* in *W*(*S*). Having finished the traversal, the computed final states of the automata correspond to their actual states in the underlying graph *G*(*S,k*)—which had not been built explicitly. Cuttlefish then characterizes the flanking vertices of the maximal unitigs in another implicit traversal over *G*(*S,k*), using the states information of the automata computed in the preceding step, and extracts the maximal unitigs on the fly.

Cuttlefish(*S*)

1. *K*← Extract-Unique-k-mers(*S*)
2. *h*← Construct-Minimal-Perfect-Hash-Function(*K*)
3. B←Compute-States(*S,h*,|*K*|)
4. **for** each *s* ∈ *S*
5. Extract-Maximal-Unitigs(*s,h,B*)

The major components of the algorithm — an efficient associative data structure for the automata and their states, computing the actual states of the automata, and extraction of the maximal unitigs from the input strings are discussed in the next subsections. Finally, the correctness of the algorithm is proven in theorem 1 in Supplementary Sec. 2.

### 4.5 Hash table structure for the automata

To maintain the (transitioning) states of the automata for the vertices in *G*(*S,k*) throughout the algorithm, some associative data structure for the vertices (i.e. canonical *k*-mers) and their states is required. We design a hash table for this purpose, with a (*k-mer, state*) key-value pairing. An efficient representation of hash tables for *k*-mers is a significant data structure problem in its own right, and some attempts even relate back to the compacted de Bruijn graph (Marchet *et al.*, 2019). In solving the subproblem efficiently for our case, we exploit the fact that the set *K* of the keys, i.e. canonical *k*-mers, are static here, and it can be built prior to computing the states. We build the set *K* efficiently from the provided set *S* of input strings using the `KMC3` algorithm (Kokot *et al.*, 2017). Afterwards, we construct a minimal perfect hash function (MPHF) *h* over the set *K*, employing the `BBHash` algorithm (Limasset *et al.*, 2017).

A perfect hash function hp for a set *K* of keys is an injective function from *K* to the set of integers, i.e. for *x*_1_, *x*_2_ ∈ *K*, if *x*_1_ = *x*_2_, then *h_p_*(*x*_1_)= *h_p_*(*x*_2_). Perfect hash functions guarantee that no hashing collisions are made for the keys in *K*. A perfect hash function is minimal if it maps *K* to the set [0, |*K*|). Since we do not need to support lookup of *k*-mers nonexistent in the input strings, i.e. no alien keys will be queried, an MPHF works correctly in this case. Also, as an MPHF produces no collisions, we do not need to store the keys with the hash table structure for collision resolution—reducing the space requirement for the structure. Although instead of the keys, we need to store the function itself with the structure, it takes much less space than the set of keys. In our setting, the function constructed using `BBHash` (Limasset *et al.*, 2017) takes ≈3.7bits/*k*-mer, irrespective of the *k*-mer size *k*. Whereas storing the keys would take 2k bits/*k*-mer.

As the hash value *h*(*x*) for some key *x* ∈ *K* is at most (|*K*| – 1), we use a linear array of size |*K*|, indexed by *h*(*x*), to store the state of the vertices (canonical *k*-mers) *x* as the hash table values. We call this array the buckets table *B*. So, the hash table structure consists of two components: an MPHF *h*, and an array B. For a canonical *k*-mer 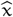, its associated bucket is in the 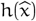’th index of B — 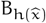 stores the state of 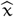 throughout the algorithm.

### 4.6 Computing the states and extracting the maximal unitigs

For a set *S* of input strings with n distinct canonical *k*-mers and an MPHF *h* over those *k*-mers, the algorithm Compute-States (*S,h,n*) computes the state of each vertex in *G*(*S,k*). Initially, it marks all the *n* vertices (i.e. their automata) with the unvisited state, in a buckets table B. Then it makes a collection of walks over *G*(*S,k*)—awalk *w* = (*v*_0_,…,*e_i_,v_i_,e*_*i*+1_,…, *v_m_*) for each *s* ∈ *S*. For each vertex *v_i_* in *w*(*s*), i.e. its associated *k*-mer instance *x* in *s*, it examines the two incident edges of *v_i_* in *w*(*s*), i.e. *e_i_* and *e*_*i*+1_, making appropriate state transition of the automaton of *v_i_* accordingly. Supplementary Fig. 1 provides a detailed overview of the DFA transition function *δ*. After the implicit complete traversal over *G*(*S,k*), the actual states of the vertices (automata) in the graph are computed correctly.

For an input string *s* ∈ *S* and the MPHF *h*, the algorithm Extract-Maximal-Unitigs(*s,h,B*) enumerates all the maximal unitigs present in *s*, using the states information of the automata present in the buckets table *B*. The enumeration is done through another implicit complete traversal over *G*(*S,k*). For a walk *w* = (*v*_0_,*e*_1_, *v*_1_,…,*e_i_,v_i_,e*_*i*+1_,…, *e_m_,v_m_*) over *G*(*S,k*) spelling *s*, say *w* enters *v_i_* through its side *s_i_*, and the class of the state of *v_i_* is *c_i_* (see Fig. 2a for state-classes). Then, *v_i_* initiates a maximal unitig traversal in *w* if:

1. *c_i_* = *multi-in multi-out;* or
2. *s_i_* is the front of *v_i_* and *c_i_* = *multi-in single-out;* or
3. *s_i_* is the back of *v_i_* and *c_i_* = *single-in multi-out;* or
4. *v*_*i*–1_ terminates a maximal unitig traversal in *w*.

*v_i_* terminates a maximal unitig traversal in *w* if:

1. *c_i_* = *multi-in multi-out;* or
2. *s_i_* is the front of *v_i_* and *c_i_* = *single-in multi-out;* or
3. *s_i_* is the back of *v_i_* and *c_i_* = *multi-in single-out*; or
4. *v*_*i*+1_ initiates a maximal unitig traversal in *w*.

The last conditions for initiation and termination do not recurse, i.e. only the first three conditions of the other case are checked for such a cross-referring condition — avoiding circular-reasoning.

Supplementary Sec. 1.2 contains the pseudo-codes of the algorithms Compute-States(*S,h,n*) and its principal component Process-*k*-mer(*x,s,h,B*), and Extract-Maximal-Unitigs (*s,h,B*).

### 4.7 Parallelization scheme

The Cuttlefish algorithm is designed to be efficiently parallelizable on a shared memory multi-core machine. The first step, i.e. the generation of the set *K* of canonical *k*-mers is parallelized in the `KMC3` algorithm (Kokot *et al.*, 2017) itself. The two major steps of it — splitting the collection of k-mers from the input strings into different bins and then sorting and compacting the bins — are highly parallelized.

The next step of constructing the MPHF with the `BBHash` algorithm (Limasset *et al.*, 2017) is also parallelized by partitioning its input key set K to multiple threads — distributing subsets of keys with some threshold size to the threads as soon as it is read off disk.

The next step of computing the states of the vertices is parallelized as follows. For each input string *s* ∈ *S, s* is partitioned into a number of uniform sized substrings. Each substring is provided to an available thread, and each thread is responsible to process the *k*-mers having their starting indices in its substring. Although the threads process disjoint sets of *k*-mer instances, they query and update the same entry in the hash table structure for identical canonical *k*-mers. Accessing the MPHF *h* concurrently from multiple threads is safe, as the accesses are read-only, whereas accessing a hash table entry itself, i.e. an entry into the buckets table B, is not, since all the threads are (read- / write-) accessing the same table B. We ensure that only a single thread can probe and / or update some specific entry at any given time through maintaining a sparse collection of access-locks into B. To ensure low memory usage, we use a bit-packed table for B. As such, even with access-locks, multiple threads may access the same underlying memory-word concurrently while accessing nearby indices. To avoid such data races, we use a thread-safe bit-packed vector (Marçais, 2020).

The last step of extracting the maximal unitigs is parallelized similarly, by distributing disjoint substrings of an input string s to different threads, where each thread is responsible to extract each maximal unitig that has its starting k-mer (in the walk spelling s) in its substring. Each maximal unitig is assigned a unique k-mer as its signature, and unique extraction of a maximal unitig is ensured by transitioning its signature k-mer to some special “output”-tagged state of the same state-class when the unitig is extracted first.

### 4.8 Asymptotics

Given a collection of strings *S*, let *m* be the total length of the input strings, and *n* be the number of distinct *k*-mers in the input. Then the running time of Cuttlefish is loosely bounded by *O*((*m* + *n*)*H*(*k*)), where *H*(*k*) is the expected time to hash a *k*-mer by Cuttlefish, assuming that `BBHash` takes *O*(*h*) time (an expected constant) to hash a machine word of 64-bits, 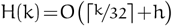. See Supplementary Sec. 3.1 for a detailed analysis. The dependence of the running time on both the input size (m) and the variation in the input (expressed through n) is exhibited for an apes dataset in Supplementary Figs. 2a, 2c, and 2b. The dependence on k is discussed at subsection 5.4, with benchmarking in Table 3.

The maximum memory usage of Cuttlefish is completely defined by the space requirement of the hash table structure. In our setting, the MPHF takes ~3.7 bits/*k*-mer. Each vertex (canonical *k*-mer) of *G*(*S,k*) canbein one of 26 different states (the state-space size for the DFA is |Q|= 26). At least ⌈log_2_(26)⌉ = 5 bits are necessary to represent such a state uniquely. Thus, the buckets table consume 5n bits in total. Therefore, the maximum memory usage of the algorithm is (8.7× *n*) = *O*(*n*) bits — translating to roughly a byte per distinct *k*-mer. This linear relationship between the memory usage and the distinct *k*-mers count is illustrated for a humans and an apes dataset at Supplementary Figs. 2b and 2d. See Supplementary Sec. 3.2 for a detailed analysis.

## 5 Results

We evaluated the performance of Cuttlefish compared to other state-of-the-art tools for constructing (colored) compacted de Bruijn graphs from whole genome references. See Supplementary Sec. 4.1 for a discussion on the definition of “color” that we adopt here. We also assessed its scaling properties, and effects of the input structure on performance. Experiments are performed on a server with an Intel Xeon CPU (E5-2699 v4) with 44 cores and clocked at 2.20 GHz, 512 GB of memory, and a 3.6 TB Toshiba MG03ACA4 HDD. The runtime and the peak resident memory usage statistics are obtained using the GNU `time` command.

### 5.1 Dataset characteristics

We use a varied collection of datasets to benchmark Cuttlefish in evaluating its performance on diverse input characteristics. First, we assess its performance on single input genomes. We use individual references of (i) a human (*Homo sapiens*, ~3.2 Gbp), (ii) a western gorilla (*Gorilla gorilla, ~3* Gbp), and (iii) a sugar pine (*Pinus lambertiana*, ~27.6 Gbp). We also evaluate its performance in building (colored) compacted de Bruijn graphs, i.e. for collections of references as inputs. We use a number of genome collections exhibiting diverse structural characteristics: (i) 62 E. Coli (*Escherichia coli*) (~310 Mbp), a dataset with small bacterial genomes; (ii) 7 Humans (*Homo sapiens*) (~21 Gbp), very closely related moderate-sized mammalian genomes from the same species; (iii) 7 Apes (*Hominoid*) (~18 Gbp), related moderate-sized mammalian genomes from the same order (*Primate*) and superfamily, but different species; and (iv) 11 Conifers (*Pinophyta*) (~204 Gbp), related huge plant genomes from the same order (*Pinales*), but different species.

The E. Coli dataset with 62 strains (available at NCBI) was used in benchmarking SplitMEM (Marcus *et al.*, 2014). The human dataset was used in benchmarking a BWT-based algorithm (Baier *et al.*, 2015) for compacted de Bruijn graph construction, and includes 5 different assemblies of the human genome (from the UCSC Genome Browser), as well as the maternal and paternal haplotype of an individual NA12878 (Utah female), from the 1000 Genomes Project. The ape dataset includes 7 available references from the orangutan and the chimp genera, a western gorilla, a human, and a bonobo. The conifer dataset consists of 9 references belonging to the pine (*Pinaceae*) family: from the pine, the spruce, and the larch genera, and a douglas fir; and 2 references from the redwoods (*Sequoioideae*) subfamily. We assembled the ape and the conifer datasets from the GenBank database (Sayers *et al.*, 2018) of NCBI.

We also assessed Cuttlefish’s performance on compacting a huge number of closely related genomes. For such, we used the 93 human references generated using the FIGG genome simulator (Killcoyne and Sol, 2014), that was used to benchmark TwoPaCo (Minkin *et al.*, 2016). Coupled with the previous 7 human genomes, this gives us a dataset of 100 human references (~322 Gbp).

### 5.2 Benchmarking comparisons

We benchmarked Cuttlefish against three other implementations of whole genome reference de Bruijn graph compaction algorithms: Bifrost (Holley and Melsted, 2020), deGSM (Guo *et al.*, 2019), and TwoPaCo (Minkin *et al.*, 2016). While TwoPaCo, like Cuttlefish, is specialized to work with reference sequences, Bifrost and deGSM are also capable of constructing the compacted graph from short-read sequences as well. We compare against Bifrost and deGSM using their appropriate settings for reference-based construction. We also note that among the tools, only Bifrost constructs the compacted de Bruijn graph without using any intermediate disk-space.

All these tools have multi-threading capability. deGSM has a max-memory option, and it is set to the best memory-usage obtained from the other tools. The rest of the tools are run without any memory restrictions. All the results reported for each tool are produced with a warm cache. Tables 1 and 2 contain a summary of the comparison results.

**Table 1.**
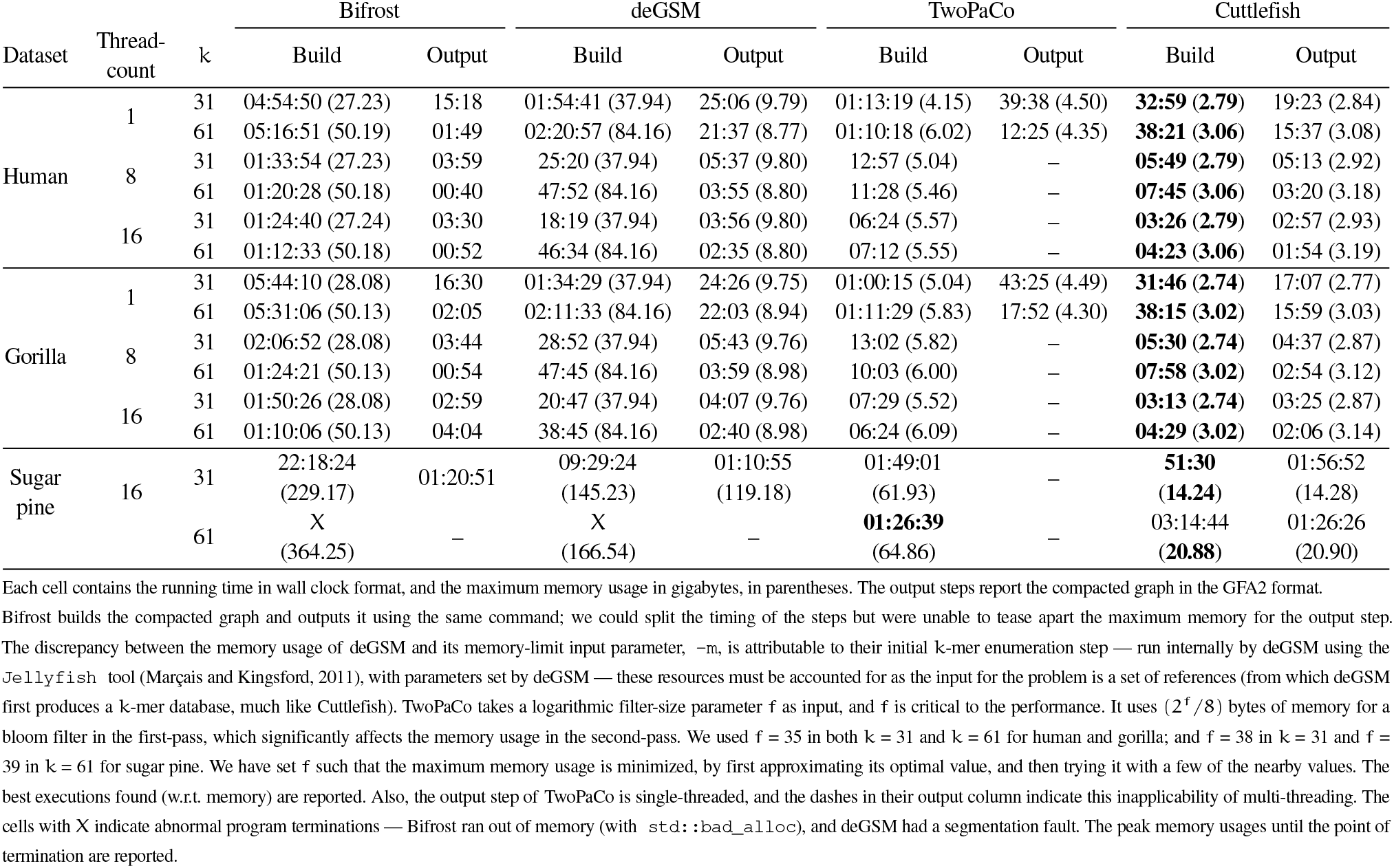
Time- and memory-performance benchmarking for compacting single input reference de Bruijn graphs.

**Table 2.** Time- and memory-performance benchmarking for compacting colored de Bruijn graphs (i.e. multiple input references) for k = 31, using 16 threads.

| Dataset | Total genome-length (bp) | Distinct k-mers count | Bifrost | deGSM | TwoPaCo | Cuttlefish |
| --- | --- | --- | --- | --- | --- | --- |
| 62 E. Coli | 310M | 24M | 1 (**0.47**) | 1 (3.34) | 1 (0.80) | 1 (0.96) |
| 7 Humans | 21G | 2.6B | 95 (29.06) | 30 (37.94) | 62 (6.14) | **21 (2.88)** |
| 7 Apes | 18G | 7.1B | 294 (100.25) | 172 (145.23) | 59 (28.87) | **25 (7.42)** |
| 11 Conifers | 204G | 82B | – | – | 981 (288.99) | **525 (84.12)** |
| 100 Humans | 322G | 28B | – | – | X (64.88) | **523 (28.75)** |

Each cell contains the running time in minutes, and the maximum memory usage in gigabytes, in parentheses. The output step is excluded from executions.

We do not compare against compaction algorithms designed for constructing the compacted de Bruijn graph only for sequencing data. These tools usually filter k-mers and alter the topology of the resulting de Bruijn graph based on certain conditions (branch length, path coverage, etc.), and these filters cannot always be finely tuned. Thus, even if ideally configured, these approaches will produce (potentially non-trivially) different compacted graphs. More importantly, however, methods that explicitly build the compacted de Bruijn graph on reference sequences can adopt or even center their algorithms around certain optimizations that are not available to methods building the compacted de Bruijn graph from reads — it therefore seems most fair not to compare methods that do not explicitly support the compacted de Bruijn graph construction from references against methods that explicitly support or are explicitly designed for this purpose. This is evident when evaluating methods that construct compacted de Bruijn graphs from reference sequences for other purposes (Minkin and Medvedev, 2020). In such cases, even authors who have worked on both state-of-the-art reference-based (Minkin *et al.*, 2016) and read-based (Chikhi *et al.*, 2016) de Bruijn graph construction methods select the former over the latter.

We verified the correctness of the produced unitigs by designing a validator, that confirms: (i) the unique existence of each k-mer in the output unitigs collection (the set of maximal unitigs is unique and forms a node decomposition of the graph (Chikhi *et al.*, 2016)); and (ii) the complete spelling-ability of the set of input references by the unitigs (i.e. each input reference is correctly constructible from the compacted graph).

A direct correspondence between the outputs of the different tools is not straightforward. See Supplementary Sec. 4.2 for a detailed discussion. The command line configurations for executing the tools, and the dataset sources are present in Supplementary Sec. 5.

### 5.3 Parallel scalability

To assess the scalability of Cuttlefish across varying number of processor-threads, we used the human genome and set k = 31, and executed Cuttlefish using 1–32 processor threads. Figs. 3a and 3b show the scaling performance charts.

**Fig. 3:**
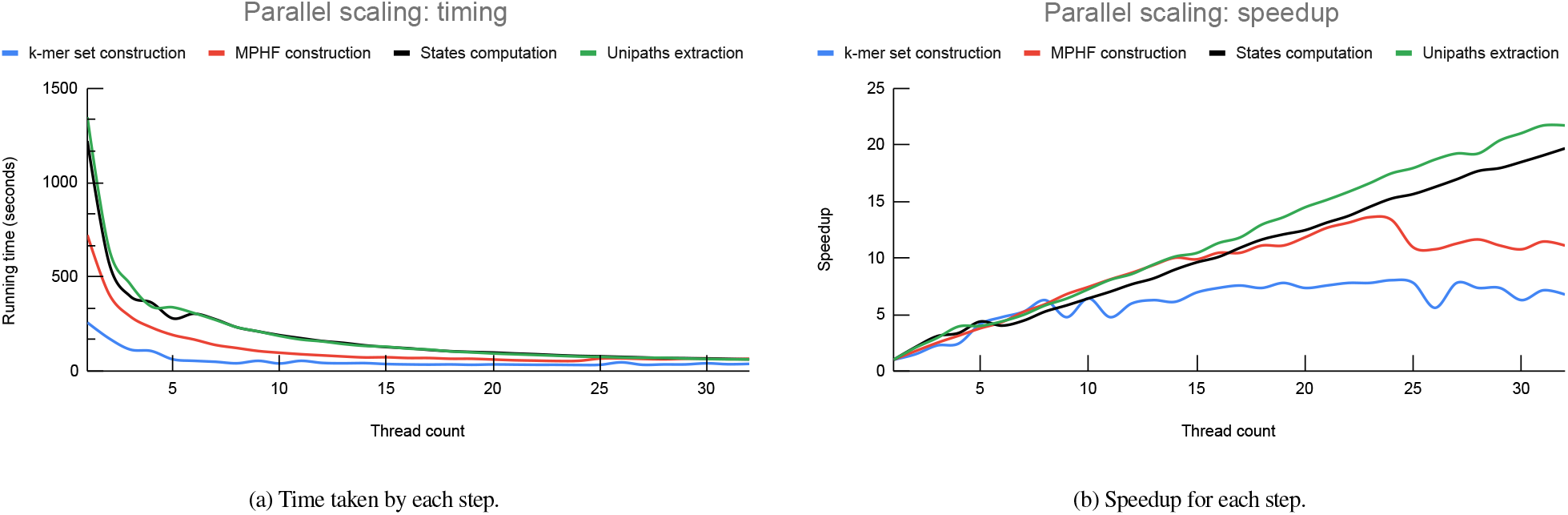
Scalability metrics of Cuttlefish for varying number of threads, using *k* = 31 for the human genome.

Generation of the set of keys (i.e. *k*-mers) is very fast for individual references, and the timing is quite low to begin with — the speedup is almost linear up-to around 8 threads, and then saturates afterwards. The MPHF (minimal perfect hash function) construction does not scale past 24 threads; which we perceive as due to limitations involving bottlenecks in disk-access throughput, and memory access bottlenecks associated to the increasing cache misses in the BBHash algorithm (Limasset *et al.*, 2017). The next two steps: computing the states of the vertices and extracting the maximal unitigs, both scale roughly linearly all the way up-to 32 threads. For the extraction step, we chose to report each maximal unitig, and skipped outputting the edges for the compacted graph in some GFA format.

### 5.4 Scaling with *k*

Next, we assess Cuttlefish’s performance with a range of different k-values. We use the 7 human genomes dataset for this experiment, and use 16 threads during construction. Table 3 shows the performance of each step with varying *k*-values.

**Table 3.** Time- and memory-performance benchmarking for the steps of Cuttlefish on the 7 human genomes dataset, across different *k*.

| $k$ | Distinct $k$-mers count (B) | Build steps (s) | | | Build Time (s) | Build memory (GB) | Output step (s) | | Output memory (GB) |
| --- | --- | --- | --- | --- | --- | --- | --- | --- | --- |
|  |  | k-mer set construction | MPHF construction | States computation |  |  | Unipaths only | GFA2 |  |
| 23 | 2.39 | 154 | 62 | 762 | 978 | 2.67 | 744 | 1345 | 2.82 |
| 31 | 2.59 | 391 | 70 | 791 | 1252 | 2.88 | 737 | 1203 | 3.01 |
| 61 | 2.96 | 439 | 200 | 797 | 1436 | 3.25 | 798 | 831 | 3.37 |
| 91 | 3.12 | 1118 | 311 | 830 | 2259 | 3.42 | 806 | 860 | 3.49 |
| 121 | 3.24 | 1483 | 902 | 841 | 3226 | 3.55 | 850 | 820 | 3.62 |

The running times are in seconds, and the maximum memory usages are in gigabytes.

We observe that the *k*-mer set construction step slows down with the increasing *k*, which may be attributable to overhead associated with the KMC3 algorithm (Kokot *et al.*, 2017) in processing larger *k*-mer sizes. Although the count of distinct k-mers grows slowly with increasing k, the MPHF construction slows down. Since the MPHF is constructed by iterating over the k-mer set on disk, the decreased speed in this step is mostly related to disk-access throughput. The steps involving graph traversals, computing the vertices’ states and extracting the maximal unitigs, are affected little in their running time by altering the value of *k*. Outputting the compacted graph in the GFA2 (and in GFA1) format takes much longer than outputting only the unitigs for lower values of k, due to structural properties of the compacted graph (see Supplementary Sec. 4.3 for a discussion).

The memory usage by Cuttlefish is directly proportional to the distinct *k*-mers-count *n*, which typically increases with *k*. Thus the maximum memory usage also grows with *k*.

Referring back to Sec. 4.8, the running time of Cuttlefish is 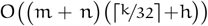, and its memory usage is *O*(8.7×*n*). Table 3 demonstrates this asymptotic increase in running time with *k* (also with effects from *n*), and the increase in memory usage with *n* (which typically grows with *k*).

### 5.5 Effects of the input structure

Next, we evaluate the effects of some of the structural characteristics of the input genomes on the performance of Cuttlefish. Specifically, we assess the impact of the genome sizes (total reference length, *m*) and their structural variations (through distinct *k*-mers count, *n*) on the time and memory consumptions of Cuttlefish. The running time is *O*((*m*+*n*)*H*(*k*)) and the memory usage is *O*(8.7 ×*n*) (see section. 4.8). The input size determines the total number of *k*-mers to be processed at the *k*-mer set construction, vertices’ states computation, and the maximal unitigs extraction steps. The variation in the input determines the performance of the MPHF construction, and indirectly affects the states computation and the unitigs extraction steps through their use of the hash table structure. We use the 7 humans and the 7 apes datasets with *k* = 31 for such, using 16 processor threads. References are incrementally added to the input set, and the performances are measured for each input instance. The benchmarks include both the steps of building the compacted graph and extracting the maximal unitigs, and are illustrated in Supplementary Fig. 2.

We observe that the running time varies both with the input size and with the structural variation. For the humans dataset, the distinct *k*-mers count does not increase much from one reference to seven: the increase is just ~5%. This is because the dataset contains references from the same species, hence the genomes are very closely related. Thus the effect of the distinct *k*-mers count remains roughly similar on all the instances of the dataset. The increase in running time is almost completely dominated by the total workload, i.e. the total size of the genomes. For the apes dataset however, the genomes are not from the same species, and the variations in the genomes contribute to increasing the distinct k-mers count to ~294% from one reference to seven. Thus the running time on this dataset increases with the total genome size, with additional non-trivial effects from the varying *k*-mers count.

On the other hand, the memory usage by the algorithm is completely determined by the distinct k-mers count, with no effect from the input size. As described at section 4.8, the memory usage of the algorithm is constant for a given dataset and k, taking ~8.7 bits/k-mer. Figs. 2b and 2d (see Supplementary Sec. 4.4) conforms to the theory—the shapes of the memory consumption and the distinct k-mers count plots are identical.

## 6 Conclusion

We present a highly scalable and very low-memory algorithm, Cuttlefish, to construct the (colored) compacted de Bruijn graph for collections of whole genome references. It models each vertex of the original graph as a deterministic finite-state automaton; and without building the original graph, it correctly determines the state of each automaton. These states characterize the vertices of the graph that flank the maximal unitigs (non-branching paths), allowing efficient extraction of those unitigs. The algorithm is very scalable in terms of time, and it uses a small and constant number of bits per distinct *k*-mer in the dataset.

Besides being efficient for medium and large-scale genomes, e.g. common mammalian genomes, the algorithm is highly applicable for huge collections of (very) large genomes. For example, Cuttlefish constructed the compacted de Bruijn graph for 100 closely related human genomes of total length ~322 Gbp in less than 9 hours, taking just 29 GB of memory. For 11 conifer plants that are from the same taxonomic order, each with very large individual genomes and having a total genome length of ~204 Gbp, Cuttlefish constructed the compacted graph in less than 9 hours, using 84 GB of memory. The next best method required more than 16 hours, using 289 GB of memory.

The improvement in performance over the state-of-the-art tools stems from the novel modeling of the graph vertices as deterministic finite-state automata. The core data structure is a fast hash table, designed using a minimal perfect hash function to assign k-mers to indices, and abit-packed buckets table storing succinct encodings of the states of the automata. This compact data structure obtains a memory usage of 8.7 bits/k-mer, leading the algorithm to greatly outperform existing tools at scale in terms of memory consumption, while being equally fast if not faster. For scenarios with further memory savings requirements, the memory usage can be reduced to 8 bits/*k*-mer, trading off the speed of the hash function.

The algorithm is currently only applicable for whole genome references. The assumption of the absence of sequencing errors in these references makes complete walk traversals over the original de Bruijn graph possible without having the graph at hand. This is not the case for short-read sequences, and the sequencing errors make such implicit traversals difficult. A significant line of future research is to extend Cuttlefish to be applicable on raw sequencing data.

On repositories with large databases containing many genome references, Cuttlefish can be applied very fast in an on-demand manner, as follows. First, one can build and store a set of hash functions, each one over some class of related genome references. This consists of the first two steps of the algorithm and is a one-time procedure, containing the bottleneck part of the algorithm. Then, whenever some set of references is to be compacted, the hash function of the appropriate super-class can be loaded into memory, and the algorithm then executes only the last two steps, which are quite fast and scalable. This works correctly because the sets of keys that these hash functions are built upon are supersets of the sets of keys being used later for the construction. Cuttlefish provides the option to save and load hash functions, making the scheme feasible.

As the number of sequenced and assembled reference genomes increases, the (colored) compacted de Bruijn graph will likely continue to be an important foundational representation for comparative analysis and indexing. To this end, Cuttlefish makes a notable advancement in the number and scale of the genomes to build compacted de Bruijn graphs upon. The algorithm is fast, highly parallelizable, and very memory-frugal; and we provide a comprehensive analysis, both theoretical and experimental, of its performance. We believe that the progression will further improve the role of the de Bruijn graph in comparative genomics, computational pan-genomics, and sequence analysis pipelines; also facilitating novel biological studies — especially for large-scale genome collections that may not have been possible earlier.

## Acknowledgments

We thank Mohsen Zakeri for providing important input into the research.

## Footnotes

1 i.e. unitigs with exactly one *k*-mer

2 This shrinks the state-space for the automata. For the subwalk (*e_i_,v_i_,e*_*i*+1_), if the edges *e_i_* and *e*_*i*+1_, are to be processed independent of each other, then during each processing, only one side of *v_i_* can be seen. This requires each side to have an unvisited state independently, making the state-space size (6× 6) = 36. From lemma 1 (see Supplementary Sec. 2), the edges *e_i_* and *e*_*i*+1_, are incident to different sides of *v_i_*. Processing them simultaneously for *v_i_* thus ensures that both the sides are seen together, making one state sufficient to denote the unvisited status’ of both the sides together. This reduces the state-space size to (5×5+1)= 26.

3 *c*_1_ and *c*_2_ do not necessarily correspond to *e*_i_ and *e*_*i*+1_, in this order; the order can also be the opposite based on the side of entrance of *w* to *v_i_*.

4 Formally, this parameter denotes the set of states reachable from the initial state under certain patterns of the input symbols. As our purpose is not the acceptance of any specific input patterns, rather just to compute the final state of each automaton, we define the accept states as the entire set of possible final states.

