## Supplementary material for "Cuttlefish: Fast, parallel, and low-memory compaction of de Bruijn graphs from large-scale genome collections"

*Bioinformatics*
doi.10.1093/bioinformatics/xxxxxx

OXFORD

Sequence Analysis

### Supplementary material for “Cuttlefish: Fast, parallel, and low-memory compaction of de Bruijn graphs from large-scale genome collections”

**Jamshed Khan [1,2] and Rob Patro [1,2,*]**

[1]Department of Computer Science, University of Maryland, College Park, MD 20742, USA and
[2]Center for Bioinformatics and Computational Biology, University of Maryland, College Park, MD 20742, USA.
`{jamshed, rob}@cs.umd.edu`

*To whom correspondence should be addressed.

#### 1 Algorithm

##### 1.1 Transition function ($\delta$) of the DFA

The transition function, $\delta$, of the automata is depicted in Fig. 1.

##### 1.2 Computing the states of the vertices

Given a walk $w(s)$ over $G(S,k)$ spelling a string $s \in S$, a k-mer $x$ in $s$, an MPHF $h$ over the canonical k-mers of $S$, and a buckets table $B$, the Process-k-mer$(x,s,h,B)$ algorithm checks the two incident edges of $x$ in the walk, as well as the state of $\widehat{x}$ from $B$ using $h$ — making appropriate state transitions for $\widehat{x}$ as required. For brevity, handling of the sentinel sides are not shown, but the extension is straightforward.

For a set of strings $S$ with $n$ distinct canonical k-mers and an MPHF $h$ over those, the Compute-States$(S,h,n)$ algorithm computes the final state of each vertex of $G(S,k)$, performing all the required state transitions. Before traversing over $G(S,k)$, it initializes the buckets table $B$ with the state *unvisited* for each vertex. Then for each $s \in S$, it initiates a walk $w(s)$ on $G(S,k)$, spelling $s$. For each k-mer $x$ encountered through $w(s)$, the Process-k-mer$(x,s,h,B)$ algorithm is executed.

##### 1.3 Extracting the maximal unitigs

For a walk $w(s)$ over $G(S,k)$ spelling the string $s \in S$, an MPHF $h$ over the canonical k-mers of $S$, and a buckets table $B$ containing the actual states of each vertex in $G(S,k)$, the algorithm Extract-Maximal-Unitigs$(s,h,B)$ enumerates all the maximal unitigs present in $w(s)$. Procedures Is-Unipath-Start$(x,w)$ and Is-Unipath-End$(x,w)$ checks whether the vertex for the k-mer $x$ initiates or terminates a maximal unitig in $w$, respectively – as per the conditions delineated in Sec. 4.6 (see main text). Procedure Extract-Substring$(s,x,y)$ extracts the substring flanked by the k-mers $x$ and $y$ from the string $s$, formatted as per the end user requirement.

#### 2 Proofs

**Lemma** 1. *For a string* $s$ *and an odd integer* $k>0$, $s$ *can only be spelled by a walk* $w$ *in* $G(s,k)$ *that enters and exits each vertex* $v_i$ *through different sides of it.*

Proof. For $G(s,k)$, consider a walk $w=(v_0,e_1,v_1,...,e_i,v_i,e_{i+1},...,e_m,v_m)$ that spells $s$. Assume that $e_i$ and $e_{i+1}$ are incident to the same side of $v_i$. Without loss of generality, say that $w$ enters $v_i$ through its front side $s_{v_i f}$. Then it exits $v_i$ using the same side. There exists some $(k+1)$-mers $x$ and $y$ in $s$ that correspond to the edges $e_i$ and $e_{i+1}$, respectively. Since $w$ spells $s$ completely, $w$ would spell $\mathrm{pre}_k(x)$ from $v_{i-1}$, $\mathrm{suf}_k(x)$ (or $\mathrm{pre}_k(y)$) from $v_i$, and $\mathrm{suf}_k(y)$ from $v_{i+1}$. So, $x$ and $y$ overlap at $v_i$ with their suffix and prefix k-mers respectively, i.e. $\mathrm{suf}_k(x)=\mathrm{pre}_k(y)$ holds.

Since $w$ enters $v_i$ through the front with $e_i$ (i.e. $x$), from the definition of incidence sides of edges (see Section 3, main text), $\mathrm{suf}_k(x)=\mathrm{label}(v_i)$. And as $w$ exits $v_i$ through the front with $e_{i+1}$ (i.e. $y$), $\overline{\mathrm{pre}_k(y)}=\mathrm{label}(v_i)$. So we get, $\mathrm{suf}_k(x)=\overline{\mathrm{pre}_k(y)}$. Combining with $\mathrm{suf}_k(x)=\mathrm{pre}_k(y)$, we get $\mathrm{pre}_k(y)=\overline{\mathrm{pre}_k(y)}$. This implies that $y_1=\overline{y_k},...,y_i=\overline{y_{k-i+1}},...,y_k=\overline{y_1}$. As $k$ is odd, the $(k+1)/2$'th character exists in $y$, and for $i=(k+1)/2$, $y_i=\overline{y_i}$ holds. But a character cannot be its own nucleotide complement, resulting into a contradiction.

Therefore, the initial assumption of $e_i$ and $e_{i+1}$ to be coincident to the same side of $v_i$ is false. Thus, $e_i$ and $e_{i+1}$ must be incident to different sides of $v_i$ for $w$ to spell $s$. ■

**Lemma** 2. *For a string* $s$ *and an integer* $k>0$, *a complete walk traversal* $w(s)$ *over* $G(s,k)$ *contains each maximal unitig of* $G(s,k)$ *as some subpath.*

Proof. Consider a maximal unitig $p=(v_0,e_1,v_1,...,e_m,v_m)$ in $G(s,k)$. A complete walk traversal $w(s)$ over $G(s,k)$ is obtained through a scan over $s$. From the definition of edge-centric de Bruijn graphs (see Section 3, main text), $w(s)$ traverses all the vertices and edges of the graph. Since $p$ is a unitig, each of its internal vertices $v_i\,(0<i<m)$ has exactly one edge on each side, and these edges chain from $v_0$ to $v_m$ to form $p$. Hence, $w(s)$ can not visit any internal vertex of $p$ without coming from $v_0$ (or $v_m$). $w(s)$ must traverse $v_0$ and then $e_1$ (or, $v_m$ and then $e_m$) at some point. From that point onwards, $p$ must be traversed entirely by $w(s)$, as it not possible to branch off from some $v_i$ without exhausting $p$. Thus, each maximal unitig $p$ is contained in $w(s)$. ■

**Lemma** 3. *For some string* $s$ *and an integer* $k>0$, *the* $(k+1)$*-mers* $e$ *and* $\overline{e}$ *correspond to the same edge in* $G(s,k)$.

Proof. Consider a $(k+1)$-mer $e$ in $G(s,k)$. $e$ can be expressed as $e=x\cdot n_x=n_y\cdot y$, where $x=\mathrm{pre}_k(e)$, $y=\mathrm{suf}_k(e)$, $n_x=e[k+1]$, and $n_y=e[1]$.

As $e=x\odot y$, $e$ induces an edge between the vertices corresponding to $\widehat{x}$ and $\widehat{y}$. From the definition of incidence sides of edges (see Section 3, main text), it is incident to the back of $\widehat{x}$ if $x=\widehat{x}$ holds, and is incident to the front of $\widehat{x}$ if $\overline{x}=\widehat{x}$ holds otherwise.

$e$'s reverse complement is $\overline{e}=\overline{n_x}\cdot\overline{x}=\overline{y}\cdot\overline{n_y}$. As $\overline{e}=\overline{y}\odot\overline{x}$, it induces an edge between the same pair of vertices $\widehat{y}$ and $\widehat{x}$. Note the order of the k-mers in $\overline{e}$. The edge is incident to the front of $\widehat{x}$ if $\overline{x}=\widehat{x}$ holds, or to the back of $\widehat{x}$ if $x=\widehat{x}$ holds otherwise.

So, $e$ and $\overline{e}$ induce edges between the same pair of vertices $\widehat{x}$ and $\widehat{y}$, and both the edges are incident to the same side of $\widehat{x}$. Similarly, it can be proved that both the edges are incident to the same side of $\widehat{y}$. Therefore, these edges are actually the same. Thus, $e$ and $\overline{e}$ induce the same edge in $G(s,k)$. ■

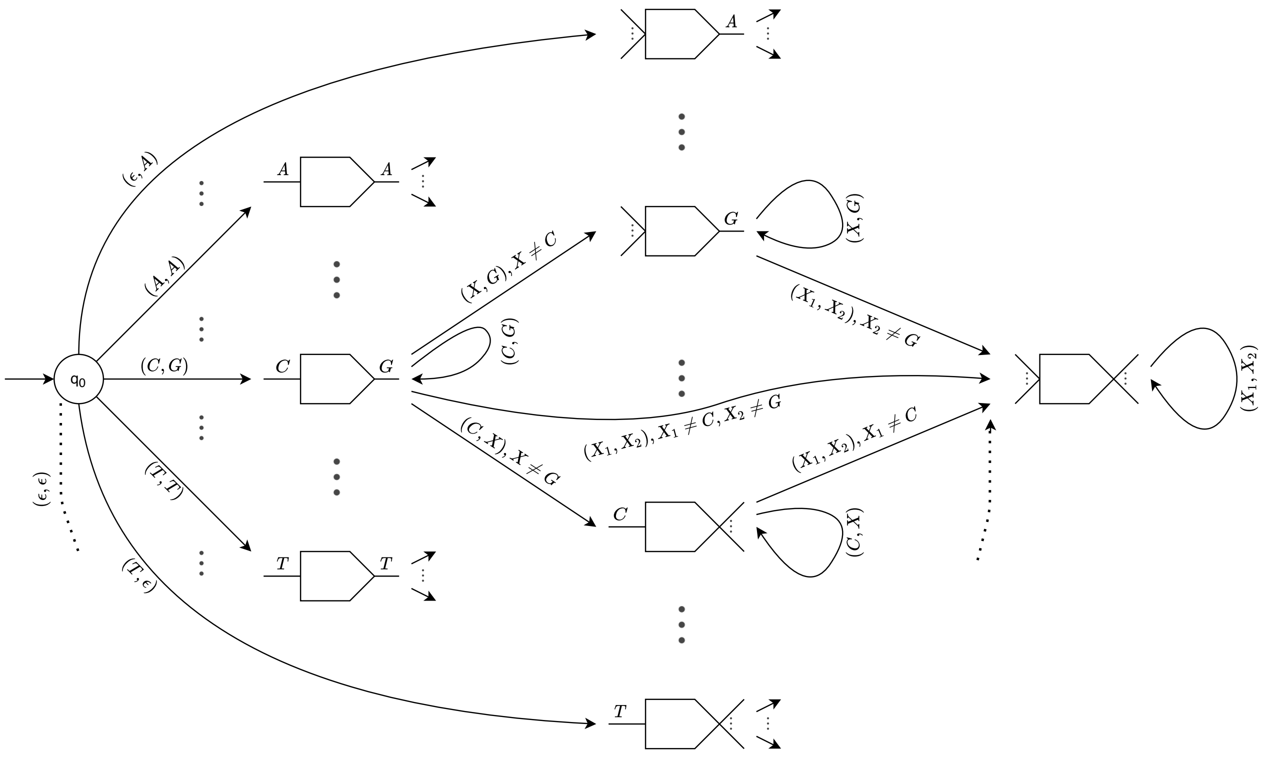

Fig. 1: Partial state-transition diagram for the automaton of a vertex. $q_0$ is the initial state, denoting the *unvisited* state. The rest 25 states are represented with the pictorial shapes as described in Fig. 2a (see main text), and we present some of the states as representatives. The $\epsilon$ characters in the input symbols denote sentinel occurrences in the corresponding sides.
For example, in a complete walk over $G(S,k)$, say an unvisited vertex $v_i$ is encountered for the first time in the subwalk $(..,e_i,v_i,e_{i+1},..)$, and the edges incident to the front and to the back of $v_i$ among $e_i$ and $e_{i+1}$ are encoded with the characters C and G, respectively. Then $v_i$ transitions from the state $q_0$ to a state of the class *single-in single-out* with the $(C,G)$ configuration. Then when $v_i$ is encountered next, its state is transitioned as per the rules delineated in Fig. 2b (see main text). Say this next occurrence of $v_i$ has edges encoded by A and G at its front and back, respectively. Then $v_i$ transitions to a state of the class *multi-in single-out* with the configuration $(X,G)$. The X here means that there are $\neq 1$ distinct edges incident to the front, and as such we do not care about what the actual edges are — thus, we are throwing away the information of the existence of the C-encoded edge at the front of $v_i$. Then as long as the next occurrences of $v_i$ in the walk has G-encoded edges at its back, irrespective of the front-incident edges, $v_i$ remains in this state. Whenever a back-incident edge is encountered with any of the other three encodings (or $\epsilon$, for a sentinel), $v_i$ transitions to the only state of the class *multi-in multi-out*, which is a dead-end for states.

**Lemma** 4. *For some string* $s$ *and an integer* $k > 0$*, a side of a vertex in* $G(s,k)$ *can have at most four distinct incident edges.*

Proof. Without loss of generality, consider the back side $s_{vb}$ of a vertex $v$ in $G(s,k)$. From the definition of incidence sides of edges (see Section 3, main text), we get that $s_{vb}$ will have an incident edge for a $(k+1)$-mer $e$ iff: (1) $\mathrm{pre}_k(e) = \mathrm{label}(v)$; or (2) $\overline{\mathrm{suf}_k(e)} = \mathrm{label}(v)$, which is the same as $\mathrm{suf}_k(e) = \overline{\mathrm{label}(v)}$.

Say that $\mathrm{label}(v) = l_v$. Recall that all our strings are over the alphabet $\Sigma = \{A,C,G,T\}$. For the first case, there can be at most $|\Sigma| = 4$ distinct edges, the corresponding set of $(k+1)$-mers being $E_1 = \{l_v \cdot c \mid c \in \Sigma\}$. Similarly, there can be at most four distinct edges for the second case, with the set of $(k+1)$-mers being $E_2 = \{c \cdot \overline{l_v} \mid c \in \Sigma\} = \{\overline{c} \cdot \overline{l_v} \mid c \in \Sigma\}$.

From lemma 3, the $(k+1)$-mers $l_v \cdot c$ and $\overline{c} \cdot \overline{l_v}$ induce the same edge for some $c \in \Sigma$. This implies that the set of edges induced from $E_1$ is the same as the set induced from $E_2$. Thus, a vertex can have at most four distinct edges incident to each side. ■

**Theorem** 1. Cuttlefish *is correct.*

Proof. Cuttlefish treats each vertex $v$ in a de Bruijn graph $G(S,k)$ as a deterministic finite-state automaton and computes its final state through a series of state-transition(s), starting from the state *unvisited*. Proving that it correctly computes the states of the vertices is trivial from the transition function $\delta$ of the automata (see Supplementary Fig. 1 for a detailed visual illustration of $\delta$). Having computed the states, it makes another set of walks over $G(S,k)$, and determines the *flanking* vertices (w.r.t. maximal unitigs) as per the conditions delineated in Section 4.6 (see main text). For an input string $s \in S$, each substring of it having terminal k-mers corresponding to such flanking vertices is reported as a maximal unitig. Since each maximal unitig of $s$ is contained in the walk over $G(S,k)$ that spells $s$ (as per lemma 2), proving that these characterized flanking vertices actually flank the underlying maximal unitigs is sufficient to prove that Cuttlefish reports only the correct maximal unitigs.

In the following, let $w = (v_0,e_1,v_1,...,e_i,v_i,e_{i+1},...,e_m,v_m)$ be a walk over $G(S,k)$ spelling some $s \in S$. Say that $w$ enters $v_i$ using its side $s_v$, and a unipath (maximal unitig) $p$ contains $v_i$.

First, we show that it is not possible for Cuttlefish to assign $v_i$ as the flanking vertex initiating $p$ when, in fact, it is not. Assume that Cuttlefish determines $v_i$ as the flanking vertex initiating $p$, whereas $v_i$ is actually not the initial vertex (in $w$) of $p$. Hence, $p$ can be extended through the side $s_v$ farther, while retaining itself a unitig. This implies that the side $s_v$ is internal to $p$. As per the definition of unitigs (see Section 3, main text), $s_v$ has exactly one incident edge $e = (u,s_u,v_i,s_v)$. Thus $w$ must enter $v_i$ using this edge $e$ from $u$. Similarly from the definition, $s_u$ also has only $e$ incident to it. Now, the first three conditions for unipath initiation state that there are $\neq 1$ edges incident to $s_v$. Therefore, Cuttlefish must have determined $v_i$ to initiate a unipath due to the last condition, i.e. it computed $u$ as a unipath terminating vertex. But similarly, the first three conditions for unipath termination state that there are $\neq 1$ edges incident to $s_u$ which is not the case. This implies that Cuttlefish determined $u$ as a unipath terminating vertex due to computing $v_i$ as a unipath initiating vertex. However, this leads to a circular reasoning, and Cuttlefish can not determine $v_i$ as a unipath initiating vertex in

```
Process-k-mer(x,s,h,B)
  x̂ ← Canonical(x)
  st ← B_h(x̂)
  if st == State(multi-in-multi-out)
      return

  entrance ← front if x == x̂; back otherwise
  e_front ← prev(x,s) if entrance is front,
            next(x,s) otherwise
  e_back ← next(x,s) if entrance is front,
           prev(x,s) otherwise
  // prev(x,s) and next(x,s) is the previous
     and the next character of x in s

  if st == unvisited
      st ← state(single-in-single-out, e_front, e_back)
  else
      if st == state(single-in-single-out, c_f, c_b)
          transition st jointly based on
          (e_front == c_f) and (e_back == c_b)
      elseif st == state(multi-in-single-out, c_b)
          transition st based on e_back == c_b
      else // st is state(single-in-multi-out, c_f)
          transition st based on e_front == c_f

  B_h(x̂) ← st
```

```
Compute-States(S,h,n)
  B ← buckets table with n entries,
      each having the unvisited state

  for each s ∈ S
      for each k-mer x in s
          Process-k-mer(x,s,h,B)
```

```
Extract-Maximal-Unitigs(s,h,B)
  start ← ϕ, end ← ϕ
  Let w be the walk spelling s

  for each k-mer x in s
      if Is-Unipath-Start(x,w)
          start ← x
      if Is-Unipath-End(x,w)
          end ← x
          Extract-Substring(s,start,end)
```

this way — resulting in a contradiction. Therefore, the initial assumption of $v_i$ being computed as a unipath initiating vertex but actually not being one is false.

Now assume that, for the walk $w$, Cuttlefish fails to determine $v_i$ as the initiating vertex for $p$, when it actually is. We proceed to demonstrate by contradiction that this can not happen. By definition, $s_v$ has $\neq 1$ incident edges as $p$ is maximal. For Cuttlefish, all four unipath initiation conditions hold false for $v_i$. Specifically, due to the first three conditions holding false, the side $s_v$ has exactly one incident edge — resulting into a contradiction. Therefore, the assumption of $v_i$ failing to be determined as a unipath initiating vertex when it actually is must be false.

Thus, Cuttlefish determines a vertex $v_i$ to be a flanking vertex initiating a maximal unitig $p$ in some walk $w$ if and only if $v_i$ actually initiates $p$ in $w$. In an analogous manner as above, it can be proven that the vertices determined as the terminating flanking vertices actually terminate maximal unitigs, and vice versa. Hence, Cuttlefish determines precisely the set of vertices that flank maximal unitigs. ■

### 3 Asymptotics

#### 3.1 Running time

Let $m$ be the total length of the input strings, and $n$ be the number of distinct k-mers in the input. While the exact asymptotic complexity for the initial phase of distinct k-mers enumeration using the `KMC3` algorithm (Kokot *et al.*, 2017) is somewhat difficult to pin down precisely, we provide a rough upper-bound for it. It consists of two major steps: bucketing the k-mers based on super k-mer signatures, and then radix sorting and compacting the buckets. The internal representation of k-mers make it possible to compare them using 64-bit machine words, i.e. 32 nucleotide-bases at a time. Splitting the k-mers collection takes time $O(\lceil k/32 \rceil m)$ (using a constant signature length), and radix sorting the buckets take total time $O(\sum_{i=1}^{b} \lceil k/32 \rceil B_i) = O(\lceil k/32 \rceil m)$, where $b$ is the number of buckets, and $B_i$ is the size of the $i$'th bucket. Therefore, a loose upper bound is $O(\lceil k/32 \rceil m)$.

Cuttlefish stores a k-mer using $\lceil k/32 \rceil$ machine words (of 64-bit). To use the minimal perfect hash function (MPHF), we convert a k-mer to a 64-bit signature, taking $O(\lceil k/32 \rceil)$ time, and use this as the key to hash. Assuming that the `BBHash` algorithm (Limasset *et al.*, 2017) takes $O(h)$ time (an expected constant) to hash each word, the time to hash a k-mer is then $H(k) = O(\lceil k/32 \rceil + h)$. `BBHash` construction algorithm requires an expected number of hashing operations linear in the number of keys $n$. Thus an expected bound for the MPHF construction time is $O(nH(k))$.

Querying and filling the buckets table in the vertices' states computation task take time $O(mH(k))$. For the next phase of the maximal unitigs extraction, the time complexity is asymptotically the same. Therefore, the (expected) total running time of Cuttlefish is $O(\lceil k/32 \rceil m + nH(k) + mH(k)) = O((m+n)H(k))$.

The dependence of the running time on both the input size ($m$) and the variation in the input (expressed through the number of distinct k-mers $n$) is exhibited for an apes dataset, detailed in Section 5.5 (see main text) and illustrated in Supplementary Figs. 2a, 2c, and 2b. The dependence on $k$ is discussed in Sec. 5.4, with benchmarking in Table 3 (see main text for both).

#### 3.2 Memory usage

The k-mer set construction by `KMC3` is done by splitting the input k-mers collection into buckets, and then sorting and compacting the buckets. This bucketing-based procedure lends effectively to operate under strict memory bounds — which can be set as deemed feasible for the working platforms.

The MPHF construction by `BBHash` is a multi-step algorithm, assigning final hash values to a subset of keys at each step and building a bit-array $H$ progressively as the output data structure. The entire k-mer set $K$ need not be present in memory, rather the algorithm processes $K$ chunk-by-chunk in each step. In the last step, the total array $H$ is present in memory; thus the algorithm requires $\Omega(|H|)$ memory. The algorithm has a parameter $\gamma$ that trades-off the construction and the query time with $|H|$, and we set it as $\gamma = 2$. This makes $|H|$ use ~3.7 bits/k-mer. Each step $d$ also requires an additional bit-array $C_d$ for the keys yet to be assigned their final hash values. If $R_d$ is the set of these keys, then $|C_d| = \gamma |R_d|$. $R_0 = K$, and $R_d$ shrinks whereas $H$ expands in size with each step. Thus, the memory usage is loosely bounded by $O(|H| + \gamma |K|)$. Specifically in our setting, the memory usage of the construction can be at most $(3.7+2) = 5.7$ bits/k-mer.

Having constructed the hash function, the buckets table $B$ is allocated as a bit-packed array. Each vertex (canonical k-mer) can be in one of 26 different states (including *unvisited*). At least $\lceil \log_2 26 \rceil = 5$ bits are necessary to represent such a state. Thus, the buckets consume $5|K|$ bits in total. Therefore, the total memory usage of the algorithm is $(8.7 \times |K|) = O(|K|)$ bits; translating to roughly a byte per distinct k-mer. This linear relationship between the memory usage and the distinct k-mers count is illustrated for a humans and an apes dataset at Figs. 2b and 2d.

### 4 Results

#### 4.1 The implied color definition in the reference de Bruijn graph

Throughout the manuscript, when we mention the colored de Bruijn graph, we refer to a very specific definition of colors. While this definition is intuitive and natural when constructing the compacted colored de Bruijn graph from a set of reference genomes, it is not the case that the Cuttlefish algorithm allows arbitrary coloring of the k-mers in the graph, at least not without another post-processing step. Specifically, in the definition adopted herein, the color set of a unitig is the subset of reference strings $s_{i_1}, s_{i_2}, \ldots, s_{i_\ell} \in S$ in which the unitig appears. This color information is implicitly encoded in the path entries of the output GFA files (the 'P' entries in GFA1 and the 'O' entries in GFA2). As a result, all unitigs produced by Cuttlefish are "monochromatic" under this coloring definition, as a change to the color set internally to a unitig would imply either a branch (which would terminate the unitig) or the start or end of some reference string and a sentinel k-mer (which would also terminate the unitig). If one were constructing the compacted colored de Bruijn graph from raw sequencing reads or from highly-fractured assemblies, then one may wish to adopt a different notion of color, wherein color sets may vary across an individual unitig. Adding such color information to the compacted de Bruijn graph produced by Cuttlefish would be possible, but would require further post-processing which we do not consider in this work.

##### 4.2 Comparison of computed maximal unitigs across algorithms

A straightforward correspondence between the outputs of the different tools is not easy. Bifrost and deGSM compact the node-centric de Bruijn graph of the references, whereas both TwoPaCo and Cuttlefish use the edge-centric variant. Also, there are two notable non-trivial differences between the graphs compacted by TwoPaCo and Cuttlefish. For TwoPaCo, each *junction* k-mer is present in all the maximal unitigs that are incident to it, so the maximal unitigs do not form a node (k-mer) decomposition of the original graph. Another difference is in the handling of inverted repeats (Sutherland and Richards, 1995) with zero intervening distance, which forms palindromic sequences. This phenomenon is surprisingly abundant relative to the expectation, e.g. large fragments of the human X and Y chromosomes are palindromic (Larionov *et al.*, 2008). An example of such is present in Fig. 1a, where the sequence GACATGTC is palindromic for the unitig GACAT. TwoPaCo canonicalizes $(k+1)$-mers instead of k-mers, and fails to capture and break such palindromic sequences, reporting the whole sequence as one unitig. Cuttlefish, however, can naturally capture these phenomena. We spot-checked that our output matches to TwoPaCo's output, when junction k-mers are properly compacted to an individual unitig (or split into their own) and when the palindromic sequences in TwoPaCo's output are broken and taken as a unitig without repeated k-mers.

##### 4.3 Scaling of GFA-formatted outputting with k

Outputting the compacted graph in the GFA2 (and in GFA1) format takes much longer than outputting only the unitigs for lower values of k. At lower k-values, the original graph contains a much larger number of maximal unitigs than it contains at higher k-values; e.g. the 7 humans dataset contains ~70M maximal unitigs at k = 23, but only ~11M at k = 121. Therefore, the compacted graph contains a much larger number of vertices at smaller k's, resulting in a much higher number of edges compared to larger k's. For example, the compacted graph for this dataset contains ~141M edges at k = 121, and ~3.5B edges at k = 23. This also increases the length (in terms of vertex count) of the paths covering the input references. So, the larger amount of disk-write operations required to output the larger GFA files makes this step relatively slower at smaller values of k, and it becomes relatively faster as k increases.

##### 4.4 Input structure effects on Cuttlefish performance

Fig. 2 demonstrates the impacts of the genome sizes (total reference length) and their structural variations (through distinct k-mers count) on the time and memory consumptions of Cuttlefish.

#### 5 Tools and datasets

For individual genome references, we used Bifrost (v1.0.5), deGSM, TwoPaCo (v0.9.4), and Cuttlefish with the following commands:

```
•Bifrost build -v -r <reference_file> \
 -o <output_file> -k <k-value> \
 -t <threads_count>
•deGSM -k <k-value> -t <threads_count> \
 -m <maximum_memory> <jellyfish_path> \
 <output_file> <reference_file>
•twopaco -o <output_file> \
 --tmpdir <working_directory> \
 -f <filter_size> -k <k-value> \
 -t <threads_count> <reference_file>
•cuttlefish build -r <reference_file> \
 -k <k-value> -s <KMC_database_prefix> \
 -t <threads_count> -w <working_directory> \
 -f <output_type> -o <output_file> --rm
```

The KMC3 database to be used by Cuttlefish is produced with the following command.

```
•kmc -k<k-value> -m<maximum_memory> -sm -fm \
 -ci0 -t<threads_count> <reference_file>
 <output_database> <working_directory>
```

For building compacted de Bruijn graphs with multiple input references, the following commands have been used:

```
•Bifrost build -v -r <reference_list> \
 -o <output_file> -k <k-value> \
 -t <threads_count>
•deGSM -k <k-value> -t <threads_count> \
 -m <maximum_memory> <jellyfish_path> \
 <output_file> <references_dir>
•twopaco -o <output_file> \
 --tmpdir <working_directory> \
 -f <filter_size> -k <k-value> \
 -t <threads_count> <ref_1> <ref_2> ... <ref_N>
•cuttlefish build -l <reference_list> \
 -k <k-value> -s <KMC_database_prefix> \
 -t <threads_count> -w <working_directory> \
 -f <output_type> -o <output_file> --rm
```

The KMC3 database is produced with the following command:

```
•kmc -k<k-value> -m<maximum_memory> -sm -fm \
 -ci0 -t<threads_count> @<reference_list> \
 <output_database> <working_directory>
```

In case of deGSM, it outputs a Burrows-Wheeler Transform (BWT) (Burrows and Wheeler, 1994) and a binary edge-sequence file, from which the GFA output has to be generated using a utility tool called ubwt (from the same authors). ubwt has been used with the following command.

```
•ubwt unipath <deGSM_BWT_file> \
 -t <threads_count> -e <deGSM_edge-seq_file> \
 -k <k-value> -o <output_file> -a G
```

Annotations of the datasets used in the benchmarks in the paper, and the URLs from which they were obtained are available at https://doi.org/10.5281/zenodo.4116608.

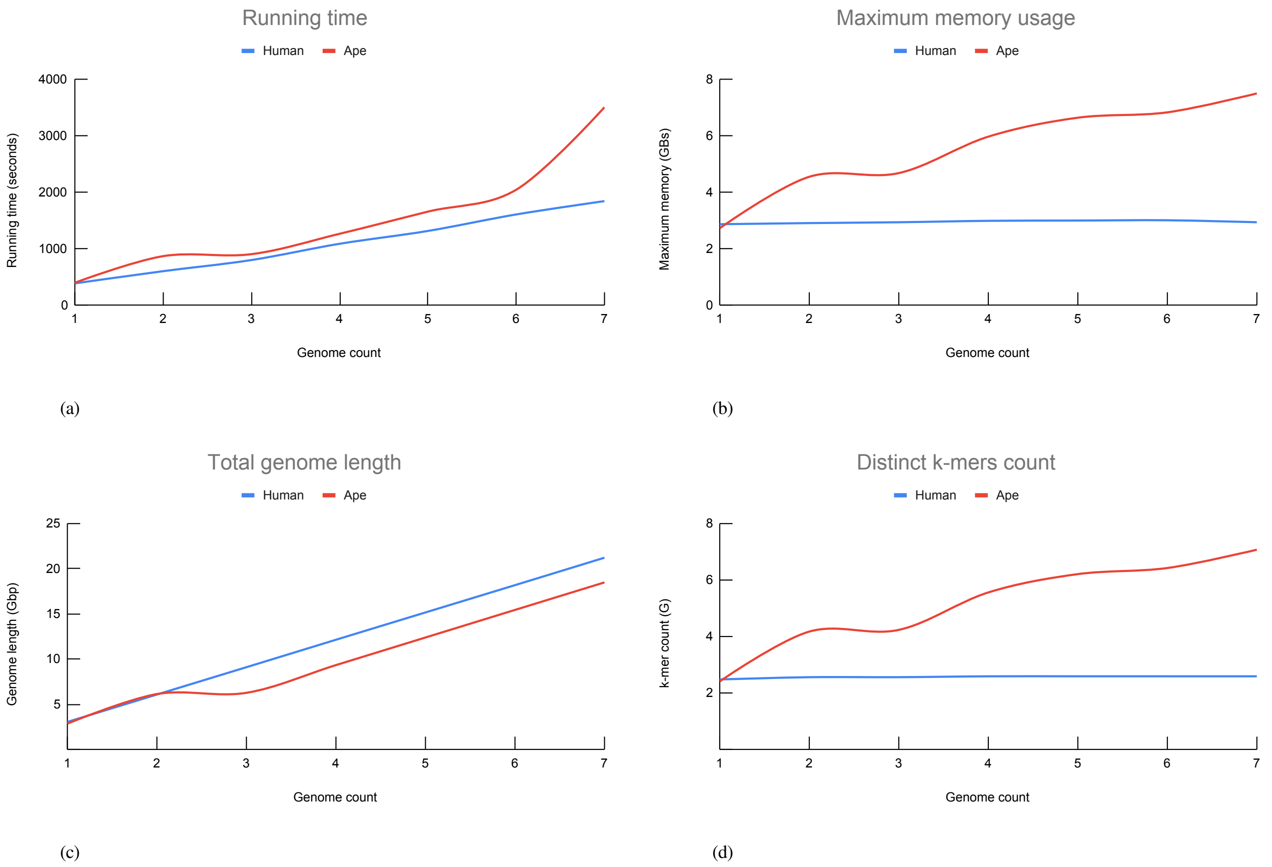


Fig. 2: Input structure effects on the performance of Cuttlefish. For genome counts varying from 1 to 7, the corresponding — (**a**) running time in seconds and the (**b**) maximum memory usage in gigabytes are reported. The corresponding (**c**) total length of the genomes and (**d**) the number distinct k-mers for each input collection are presented too.
